# Histogenesis of the reticulum of non- descript goat (*Capra hircus*) of India

**DOI:** 10.1101/2020.02.14.950311

**Authors:** Varsha Gupta, Muneer Mohhamad Farooqui, Ajay Prakash Sharma, Archana Pathak

**Author notes:** Assistant Professor, Dept. of Veterinary Anatomy. COVSc. DUVASU, Mathura. Professor, Dept. of Veterinary Anatomy. COVSc. DUVASU, Mathura. Professor and Head, Dept. of Veterinary Anatomy. COVSc. DUVASU, Mathura. Professor, Dept. of Vety. Anatomy. COVSc. DUVASU, Mathura. **Corresponding Author**: Varsha Gupta, Assistant Professor, Department of Veterinary Anatomy, College of Veterinary Sciences, DUVASU, Mathura, Uttar Pradesh, India, postal code- 281001.

## Abstract

The study was conducted on 36 developing reticulum from healthy and normal embryos/ foeti of Indian goat. Embryos/foeti were assigned into three groups. Histo-differentiation of reticulum of goat stomach took place at 38 days of gestation. The wall of foetal reticulum was made up of three strata i. e. epithelium, pleuripotent blastemic tissue and serosa up to 49 days of fetal age and definite four layers *viz.* epithelium, propria-submucosa, tunica muscularis and serosa were observed first at 51 days of gestation. Upto 100 days of gestation the reticulum was lined by undifferentiated stratified epithelium and thereafter gradually transformed into stratified squamous epithelium. Keratinization was noticed at term. Primary and secondary reticular crests made their appearance at 60 and 112 days of gestation, respectively. Corial papillae were detected in primary and secondary reticular crests at 121 and 145 days of gestation, respectively. Near term, the core of the reticular crest also contained muscularis mucosae in its proximal 1/3^rd^ region while core of the corial papillae contained only lamina propria. Reticular, collagen and elastic fibers came into sight at 46, 100 and 134 days of gestation, respectively.

**Summary statement:** The study was conducted on intrauterine reticulum of Indian goat. From this study it can be concluded that the histogenesis of reticulum was almost completed in prenatal life. However, to become functional it still required more time as the relative sizes of reticulum and process of keratinization were yet to be completed.

## 1. Introduction

Goat is important in arid, semiarid and mountainous region where crop and dairy farming are not economically feasible. The ability to browse and optimize the use of grazing land has been linked to the peculiar nature of the stomach of ruminants [1]. Reticulum plays a crucial role in the ruminant digestive tract because the primary cycle of ruminal motility always starts with reticular contraction. Documentation of normal, embryonic and fetal development is necessary to understand the consequences of harmful influences at various stages of gestation [2]. Literature on development and growth of reticulum has been carried in other ruminants; however, these research findings cannot easily be adapted to local breeds of India due to variation in genetic makeup, climate, vegetation and feeding regimen. There is a need of biological and anatomical data on development of reticulum on goat. Therefore the study was designed to get necessary knowledge of accurate day of appearance of primordium of reticulum, its differentiation and maturation.

## 3. Results

### Group I

The developing digestive tube was first observed at 32 days of gestation. At this stage of gestation the wall of digestive tube was irregularly thick (119.13±8.55µm) and encircled a narrow lumen. Stomach was consisted of three strata viz. epithelium, pleuripotent blastemic tissue and serosa (Figure 1). The epithelium was undifferentiated stratified type. Most of the cells were columnar in shape along with few cuboidal cells and contained elongated or spherical centrally placed nuclei. Their cell boundaries were indistinct and infranuclear zone was lightly eosinophilic or pale. The nuclei of cells of the upper layer were also elongated and located towards base. The cytoplasm was eosinophilic in supranuclear zone. Pleuripotent layer was thickest among the three layers and covered most of the stomach wall. Cells of the pleuripotent blastemic layer consisted of several irregularly arranged polygonal to irregular shaped mesenchymal cells with ground substance. Few blood vessels and immature red blood cells were also present. The serosa consisted of single layer of cuboidal cells present in patches. The nuclei of these cells were rounded and darkly stained.

**Figure 1.**
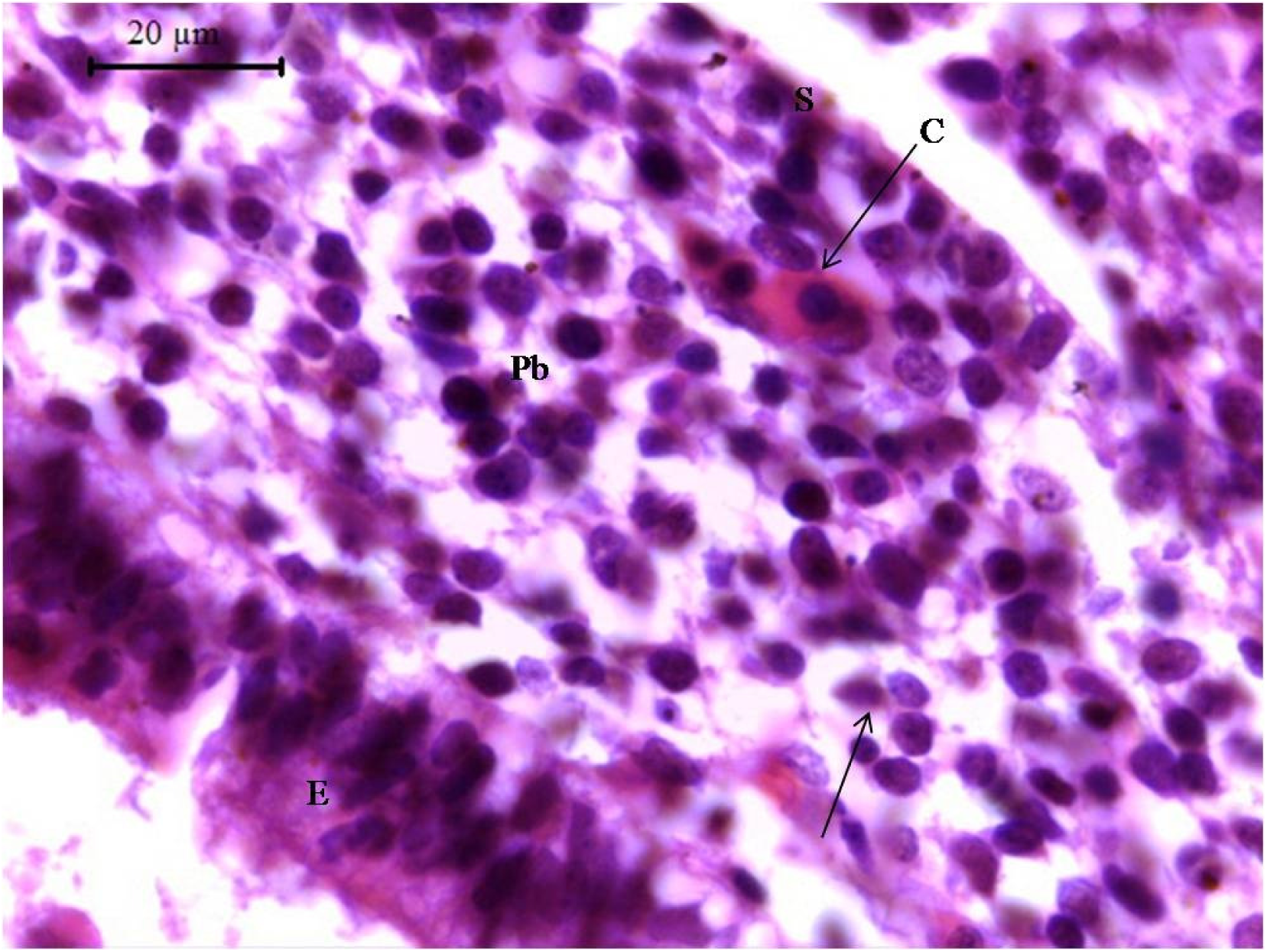
Photomicrograph of 32 day old goat foetus showing epithelium (E), pleuripotent blastemic tissue (Pb), serosa (S), capillary (C) and differentiating mesenchymal cell (arrow). H & E X 1000

Histo-differentiation of reticulum of goat stomach took place at 38 days of gestation. The wall of foetal reticulum was made up of three strata i. e. epithelium, pleuripotent blastemic tissue and serosa up to 49 days of fetal age (Figure 2). In all the foeti the reticular epithelium was divided into basal and superficial zones. The parenchyma of the developing reticulum consisted of compact cell mass in the center and degenerative process and beginning of canalization or future lumen formation was observed at 38 days of gestation from the posterior end. The epithelium was undifferentiated stratified type and two to four and four to five layered thick at 38 and 44 days of gestation, respectively (Figure 2). Basal zone had two layers upto 44 days of gestation. The cells of the basal zone were simple cuboidal shaped with indistinct cell boundaries. The vesicular nuclei were spherical in shape and had eosinophilic nucleoplasm with uniformly distributed nuclear chromatin. The cytoplasm was eosinophilic at 38 days and infranuclear zone was pale and vacuolated at 44 days of gestation. In few cells nucleoli were centrally placed. The cells of second and other layer of basal zone were either low columnar or polyhedral in shape. These cells contained vesicular, spherical or round shaped centrally placed nuclei. Their cytoplasm was either lightly eosinophilic or pale. The cells of superficial zone were arranged in one to two layers and were cuboidal or polyhedral shaped, with centrally placed spherical nuclei or without nucleus. The nuclei were either eccentric or absent at 46 days of gestation. The nucleoplasm was eosinophilic with evenly distributed nuclear chromatin. Cell boundaries were eosinophilic and cell cytoplasm was lightly eosinophilic or pale. The average thickness of the epithelium was 18.57 ± 2.97µm.

**Figure 2.**
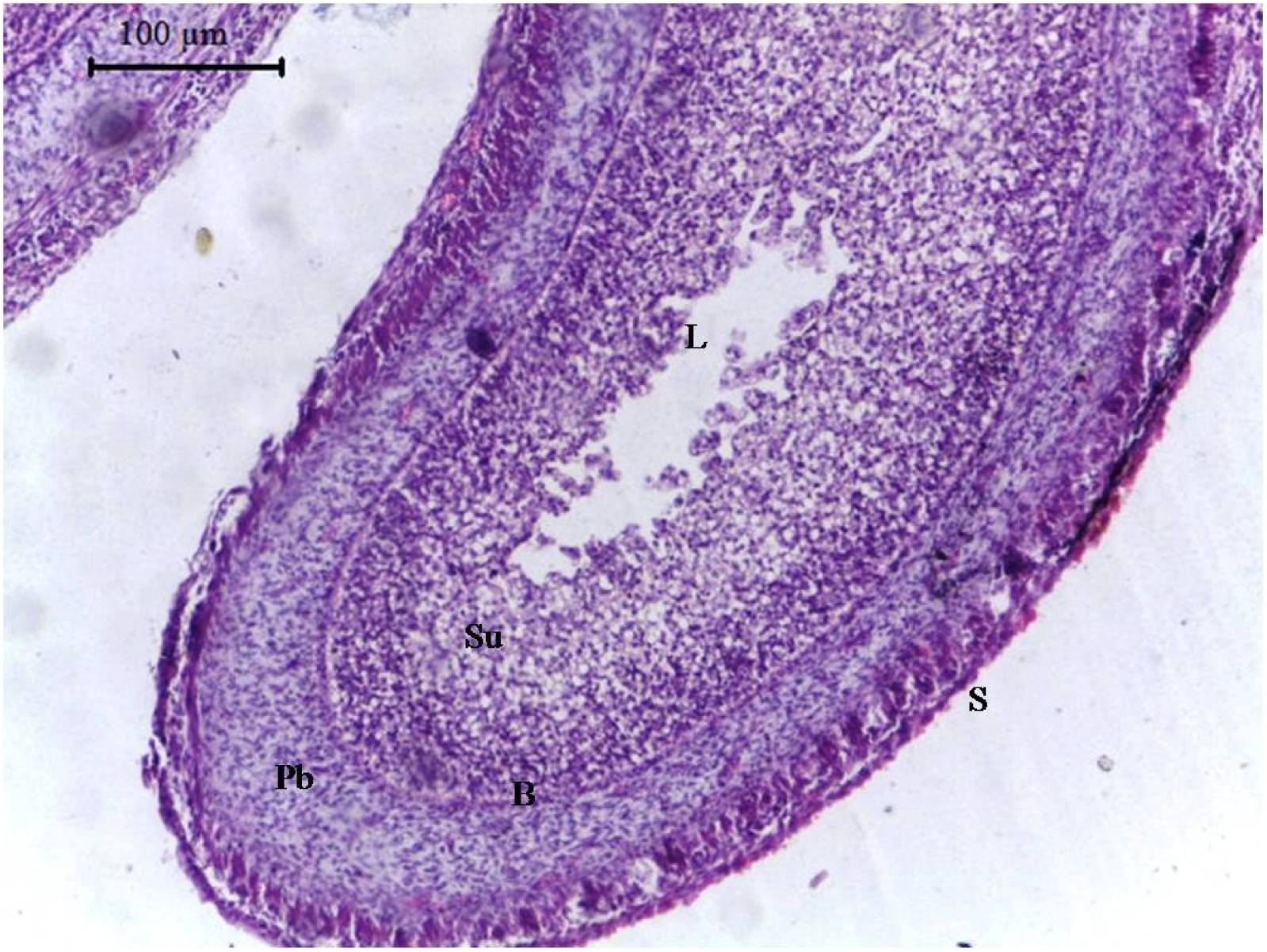
Photomicrograph of 46 day old goat foetus showing process of lumen formation (L), superficial (Su) and basal (B) zones of epithelium, pleuripotent blastemic tissue (Pb) and serosa (S).H & E X 100

The blastemic tissue was enriched with differentiating irregular mesenchymal cells with mitotic figures in few cells, differentiating fibroblasts, differentiating myocytes, capillaries, neuronal elements and ground substances (Figure 2). Clear-cut demarcation between lamina propria and submucosa was not evident in this group. Differentiating myocytes were present in 2-3 rows and most of them were loosely arranged. These cells were elongated or fusiform shaped with centrally placed elongated nuclei. Nuclear chromatins was evenly distributed and in few cells adhered to nuclear membrane. Cytoplasm of these cells was highly eosinophilic. In the close vicinity of differentiating myocytes the blastemic tissue was less cellular. Differentiating mesenchymal cells present in between these myocytes. Close to the differentiating tunica serosa loosely arranged nerve element cells were noticed. It contained two types of cells. One was large, spherical to ovoid in shape with an indistinct contour. Nuclear chromatin of these cells was evenly distributed and lightly stained. Such cells could be spoken as ganglionic cells. Another type was small with indistinct cell boundary, referred as supporting cell. Nuclei of these cells were spherical in shape with darkly stained chromatin and cytoplasm was pale. With the advancement of age myocytes were arranged in clusters of 4-5 cells and they were directed in different directions in same section. Fine isolated reticular fibrils were noticed in blastemic tissue around the differentiating myocytes and nerve elements at 46 days of gestation. The blastemic tissue was lined by a single layer of differentiating mesothelial cells with indistinct contour to form tunica serosa (10.23 ± 0.05 µm thick).

### Group II

Definite four layers of the wall viz. epithelium, propria-submucosa, tunica muscularis and tunica serosa were observed first at 51 days of gestation (Figure 3). At 51 to 60 days basal zone comprised of four to five cell layers which decreased gradually after 60 days of gestation. The cells of upper layer of basal zone were cuboidal to low columnar in shape with spherical or elongated nucleus. With the advancement of age number of layers of superficial zone increased and number of anucleated cells became abundant in this zone. The number of layers in superficial zone was 8-10, 12-15, 25-30 and 30-35 at 51, 60, 82 and 100 days of gestation, respectively. At 87 days of gestation few cells became columnar in shape and showed cytoplasmic processes at their apical end, could be referred as future stratum spinosum. Above this few polyhedral shaped cells with spherical and eccentrically placed nuclei were noticed in one or two layer. The average thickness of epithelium significantly increased from group I to II and was 132.25 ± 37.88 µm.

**Figure 3.**
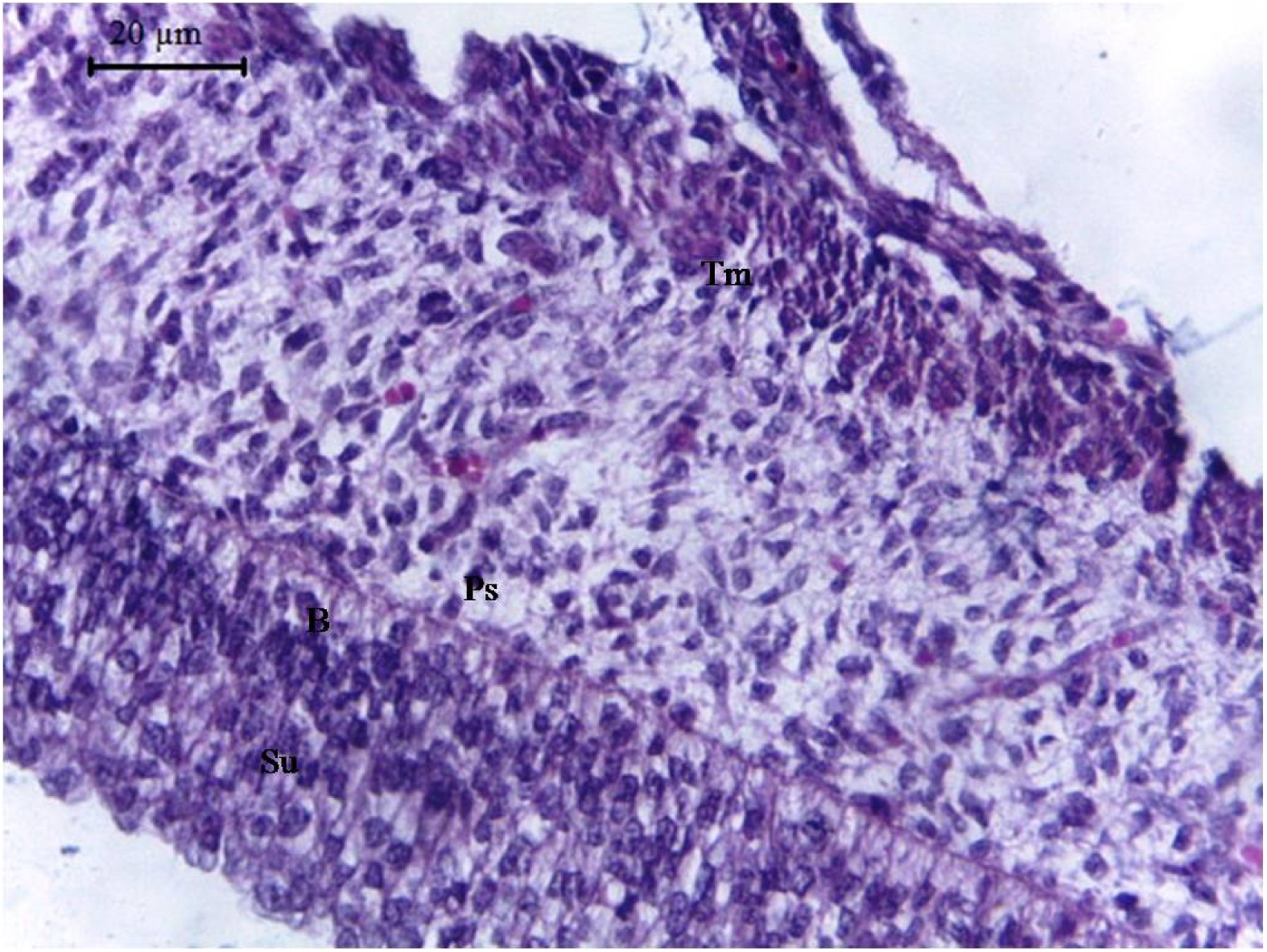
Photomicrograph of section of 51 day old goat foetal reticular wall showing superficial (Su) and basal (B) zones of undifferentiated stratified epithelium, propria submucosa (Ps) and tunica muscularis (Tm). H & E X 400

At 60 days of gestation the surface epithelium raised at certain places from its original level which may be the forerunner of the future reticular crest. At 76 days the basal layers got evaginated in the form of crecenteric shaped structure, the primary reticular crest. The crest became low pyramid at 82 and pyramidal shaped at 87 days of gestation. The core of primary reticular crest was comprised of differentiating mesenchymal cells along with capillary network below the basal layer of epithelium. The average height of the primary reticular crest was 75.62 ± 12.32 µm and width of the primary reticular crest at its origin, middle and tip was 85.75 ± 19.13, 47.56 ± 14.92 and 18.9 ± 6.08 µm, respectively. The average distance between two reticular crests was 210.91 ± 36.64 µm

At 51 days of gestation propria-submucosa and the tunica muscularis became a separate tunics but clear cut lamina muscularis could not be observed (Figure 3). The future lamina propria was darkly stained and more cellular, whereas, submucosa was lightly stained and less cellular with profound ground substance. At 70 days thin, fine, wavy isolated reticular fibrils were observed in lamina propria whereas these fibers were numerous in submucosa. These fibers became courser at 100 days of gestation and in future submucosa these fibers showed branching and anastomosing pattern at few places. Few immature isolated collagen fibers were perceived in the future submucosa at 100 days of gestation. Average thickness of propria-submucosa was 70.90 ± 14.47 µm.

Few myocytes just below the propria submucosa were arranged in clusters were noticed at 51 days of gestation (Figure 3). These clusters were of two to four cells at 70 days and four to five cells at 82 days of gestation. These clusters form small inner circular and outer longitudinal bundles. At 87 days of gestation reverse orientation of tunica muscularis was noticed as compared to 70 days (Figure 4). Differentiating mesenchymal and fibroblasts were present in between these clusters. Thin reticular fibrils were present around myocytes. These fibrils incompletely encircled clumps of myocytes and became course and numerous at 100 days of foetal age. The clusters of myocytes which were close to the submucosa were incompletely encircled by thin collagen fibrils. Around myocytes few thin collagen fibrils were also noticed. Few discontinuous collagen fibrils were also noticed in between the longitudinally arranged smooth muscle cells while they could not be observed around circularly oriented smooth muscle cells. Average thickness of tunica muscularis was 89.10 ± 32.58 µm.

**Figure 4.**
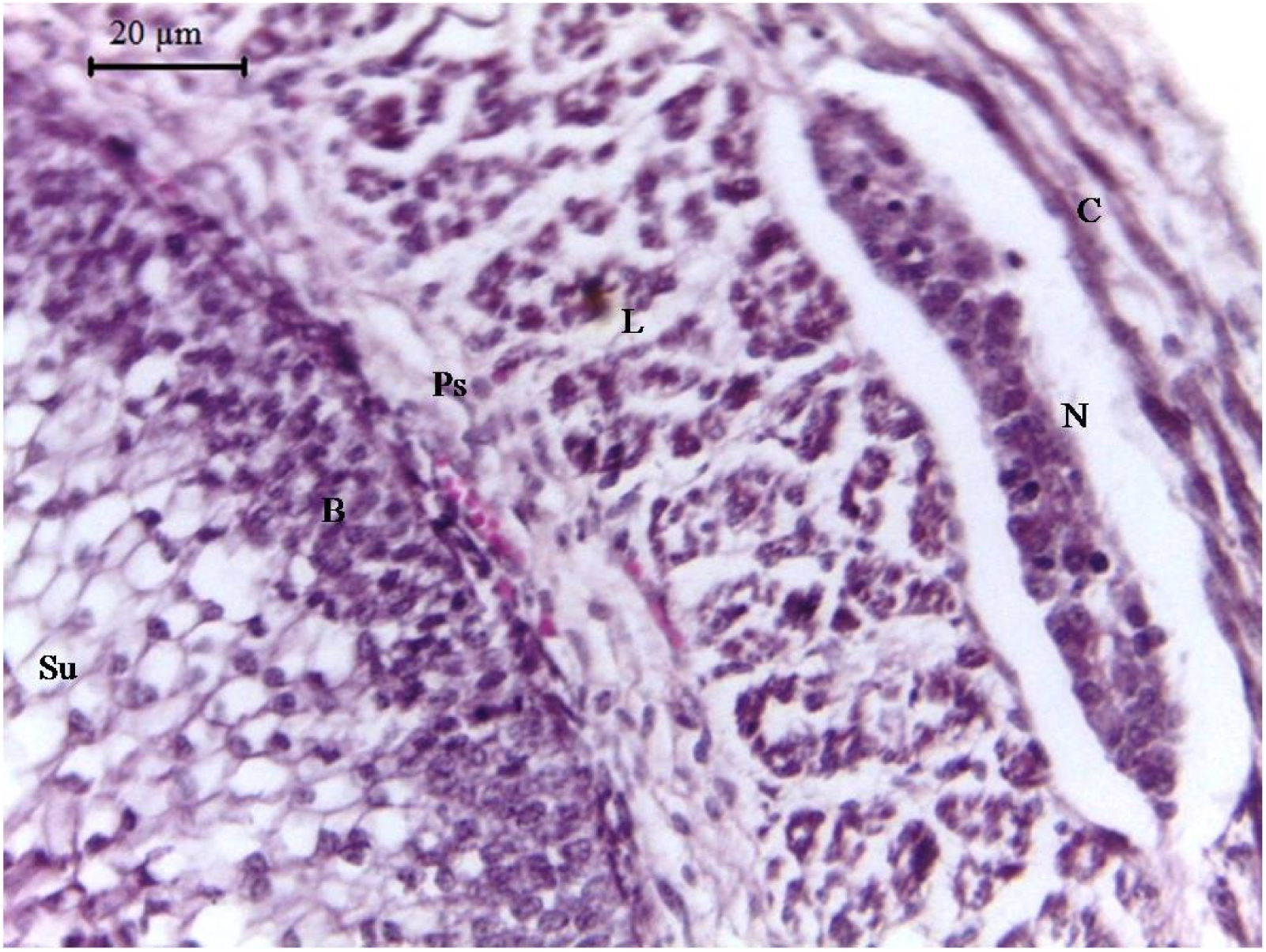
Photomicrograph of section of 87day old goat foetal reticular wall showing superficial (Su) and basal (B) zones of undifferentiated stratified epithelium, propria submucosa (Ps), Inner longitudinal (L) and outer circular (C) muscle bundles of tunica muscularis and neuronal element (N). H & E X 400

From 2^nd^ group onwards the tunica serosa was lined by a layer of flat epithelial cells. The mesothelium was well supported by loose, vascular connective tissue and had major amount of ground substance. Sporadic, fine collagen fibers were also noticed. Tunica serosa measured 12.25 ± 4.53 µm thick in group II.

### Group III

With the advancement of gestation cytological character of cells of deepest layer remained constant however, their cytoplasm became more eosinophilic and foamy at some places in full term foetal reticulum (Figure 9). The number of layers of basal zone remained constant i. e. two to three layers. At 102 days of gestation few flat cells among cuboidal cells were noticed in the topmost layer of superficial zone. At full term few cells of the top most layers became flat, had lost their nuclei and contained highly eosinophilic cytoplasm, referred as beginning of formation of stratum corneum (Figure 9). The average thickness of stratum corneum was 4 µm at the tip of papillae and 3.4 µm at the sides of papillae. The cells of reticular crest were arranged in two to three rows at the base and middle of the crest while they were in single layer at the tip of the crest. The height and inter-crest distance of the primary reticular crest increased with the advancement of age. The height of the primary reticular crest was 290.01 ± 78.01 µm. The width of the primary reticular crest at its origin, middle and tip was 93.78 ± 37.07, 38.95 ± 19.15 and 21.52 ±11.45 µm. Distance between two reticular crests was 616.29 ± 85.5 µm.

**Figure 5.**
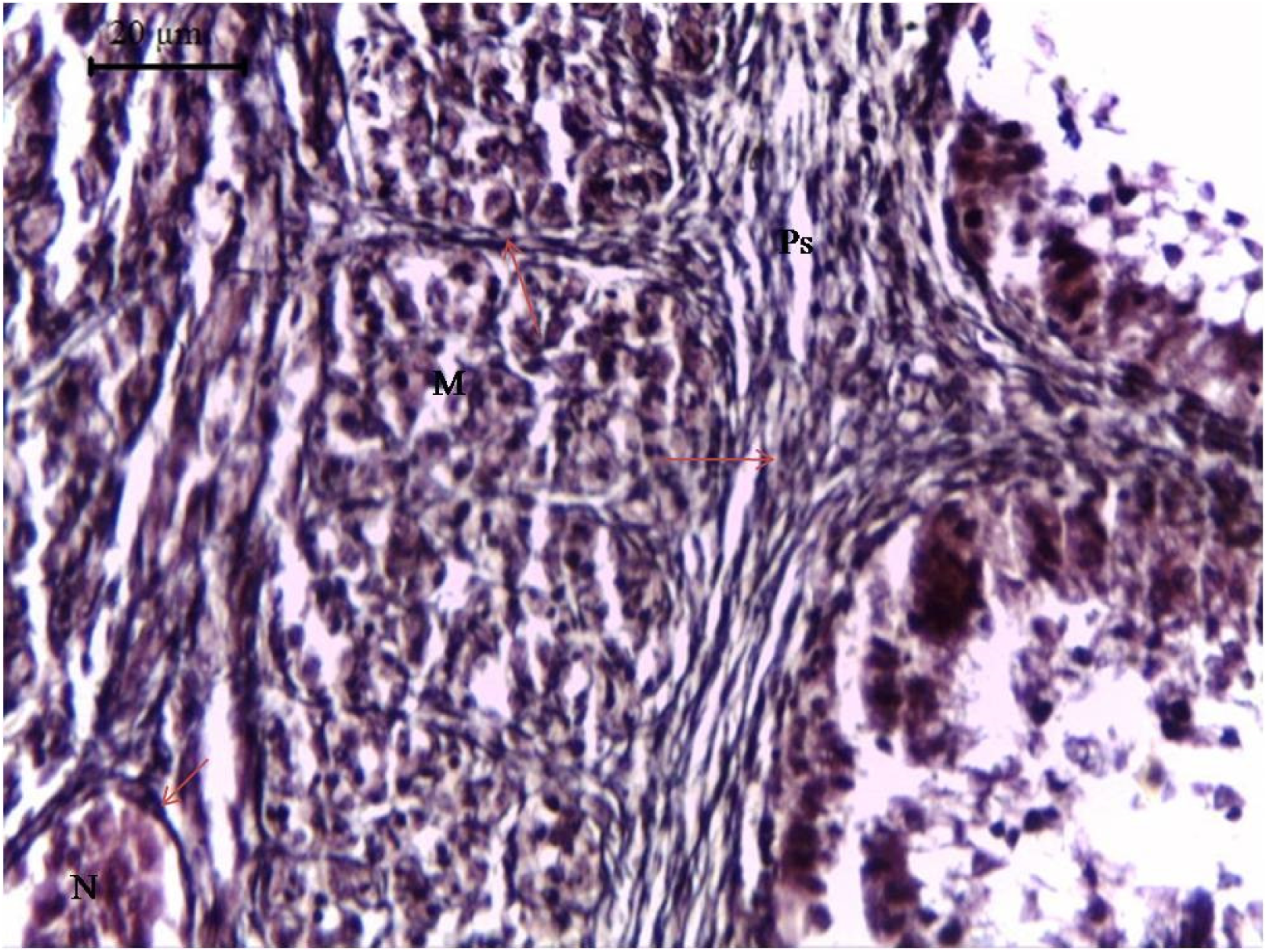
Photomicrograph of section of 118day old goat foetal reticular wall showing fine to coarse reticular fibers (arrow) in propria-submucosa (Ps) and coarse reticular fiber (arrow) in between muscle bundles (M) and around neuronal element (N). Wilder’s reticular stain X 400

**Figure 6.**
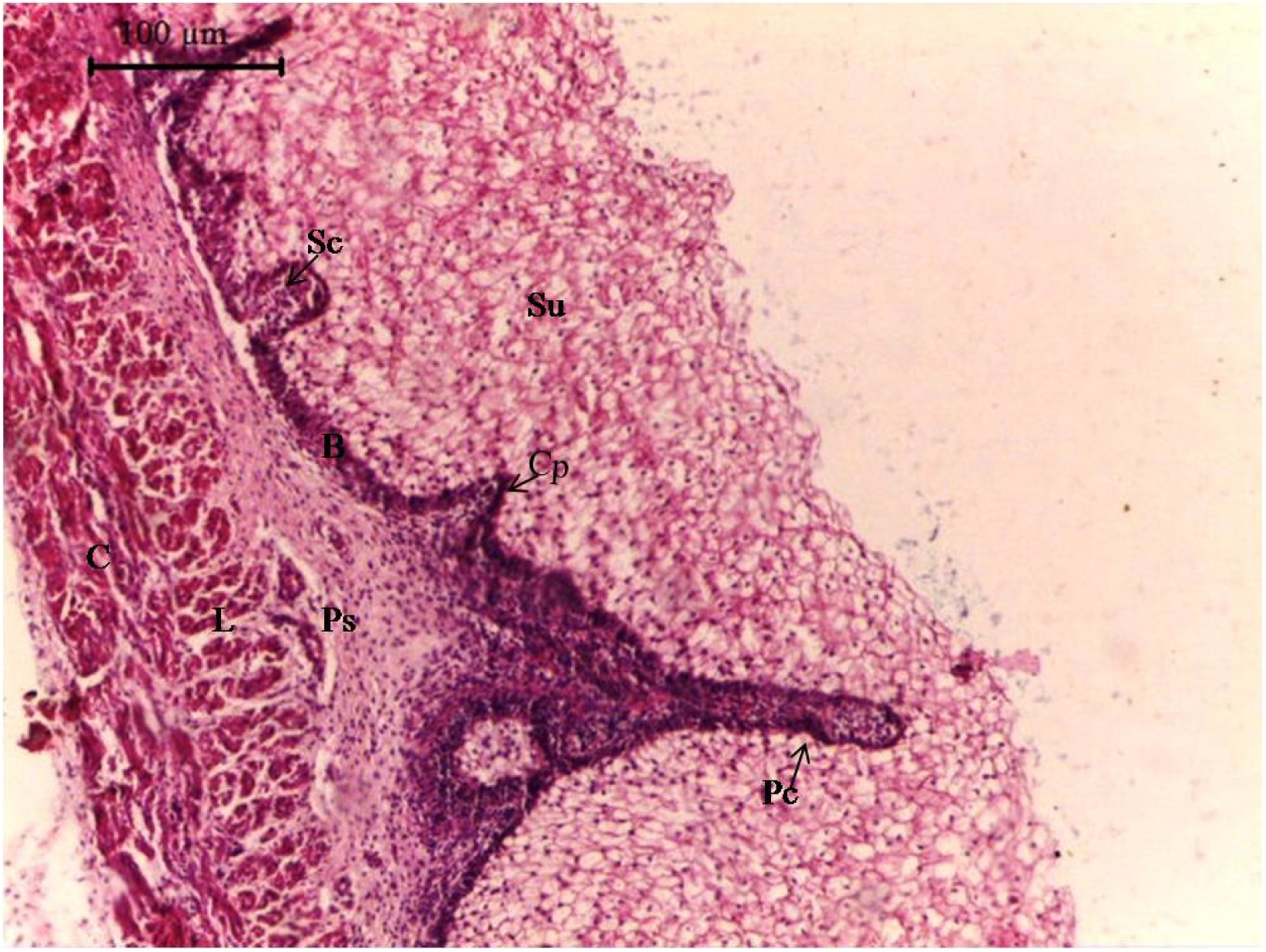
Photomicrograph of section of 121 day old goat foetal reticular wall showing superficial (Su) and basal (B) zones of epithelium, propria submucosa (Ps), Inner longitudinal muscle bundles (L), outer circular muscle bundles (C) of tunica muscularis, primary reticular crest (Pc), secondary reticular crest (Sc) and corial papilla (Cp). H & E X 100

**Figure 7.**
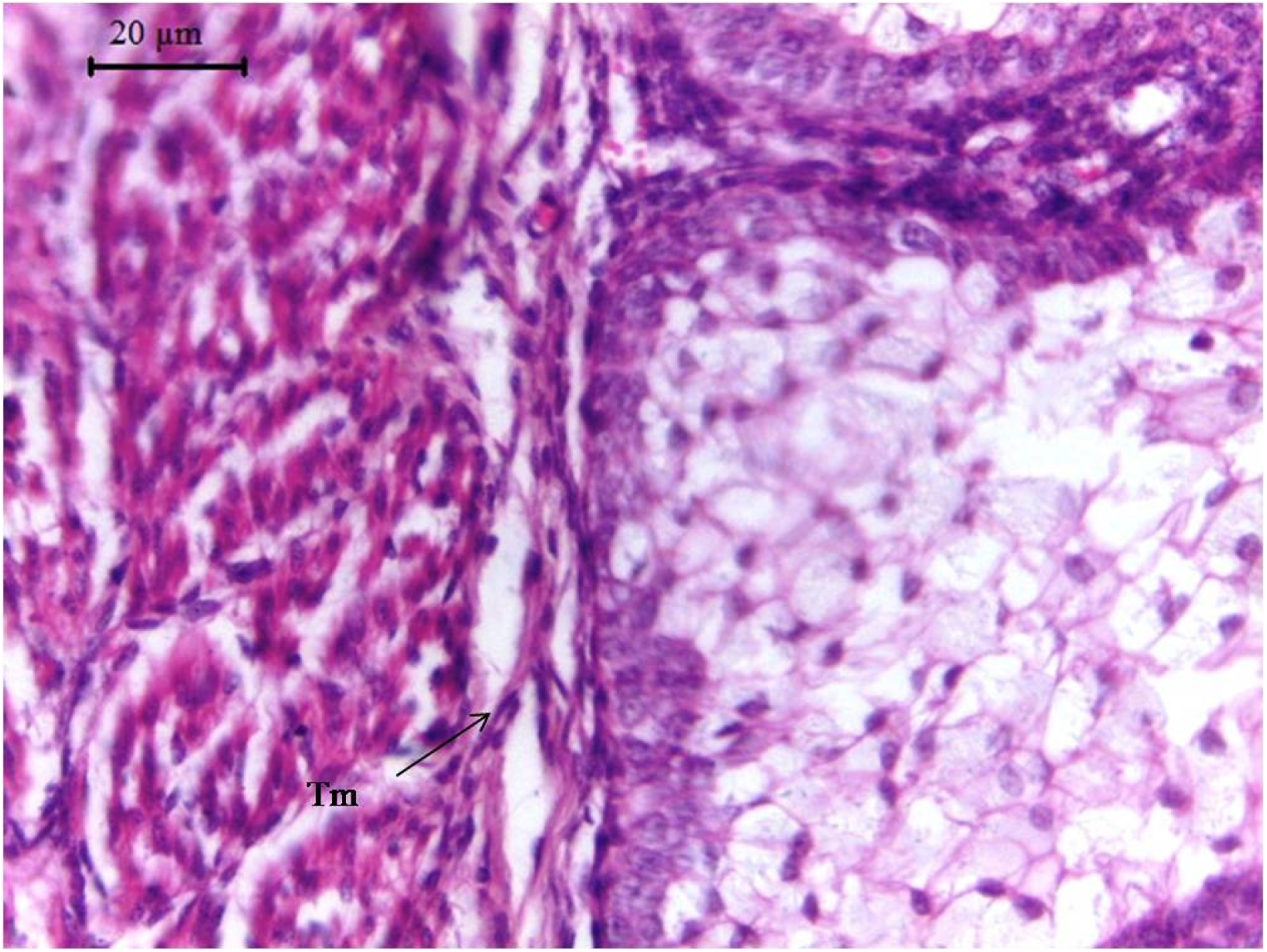
Photomicrograph of section of 134 day old goat foetal reticular wall showing delamination of smooth muscle cells from tunic muscularis (arrow) and bundles of smooth muscle cells (Tm). H & E X 400

**Figure 8.**
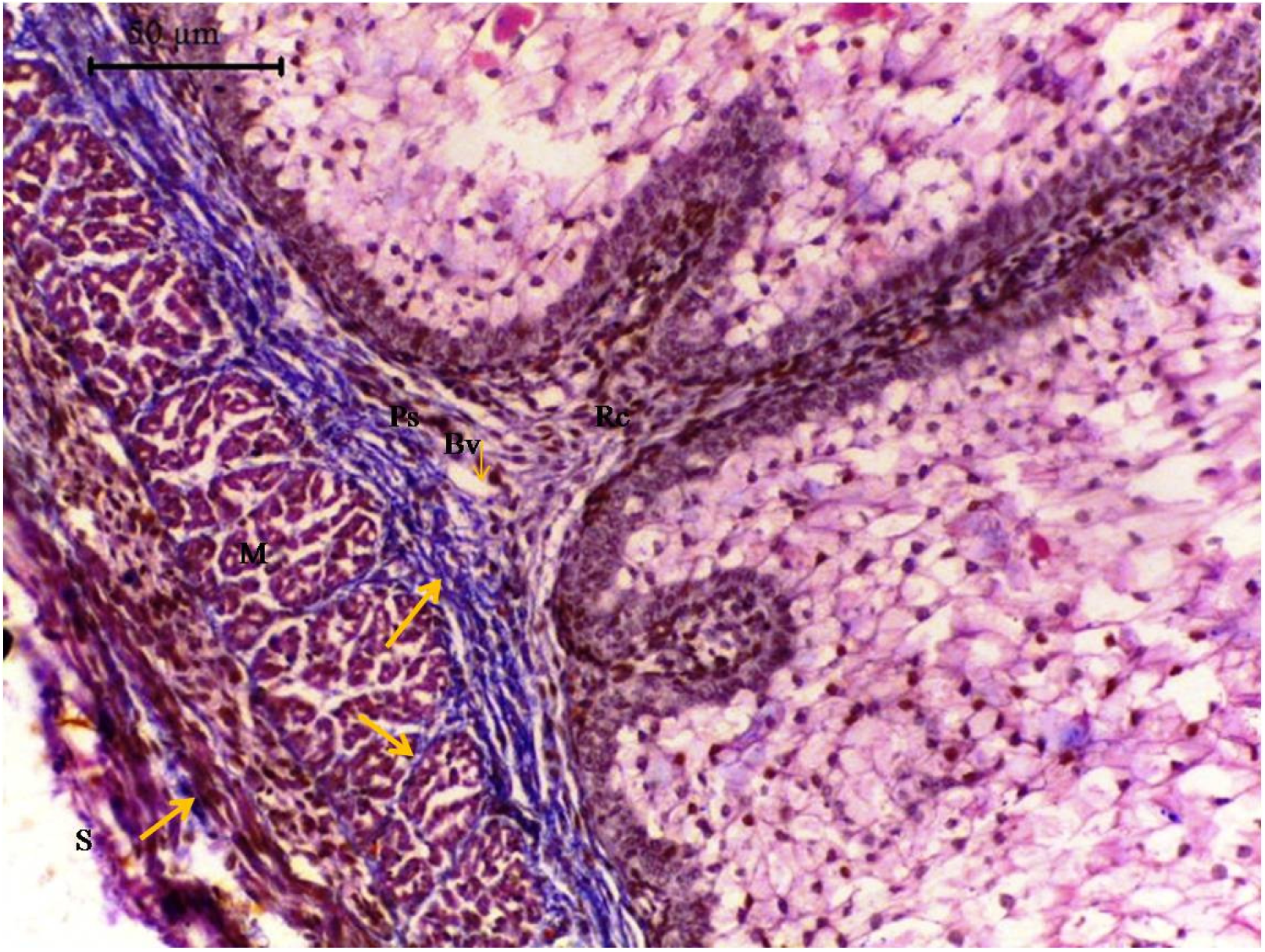
Photomicrograph of section of 134 day old goat foetal reticular wall showing coarse collagen fibers (arrow) in propria submucosa (Ps), base of the reticular crest (Rc), in between muscle bundles (M), around blood vessel (B) and serosa (S). Masson’s trichrome method X 200

**Figure 9.**
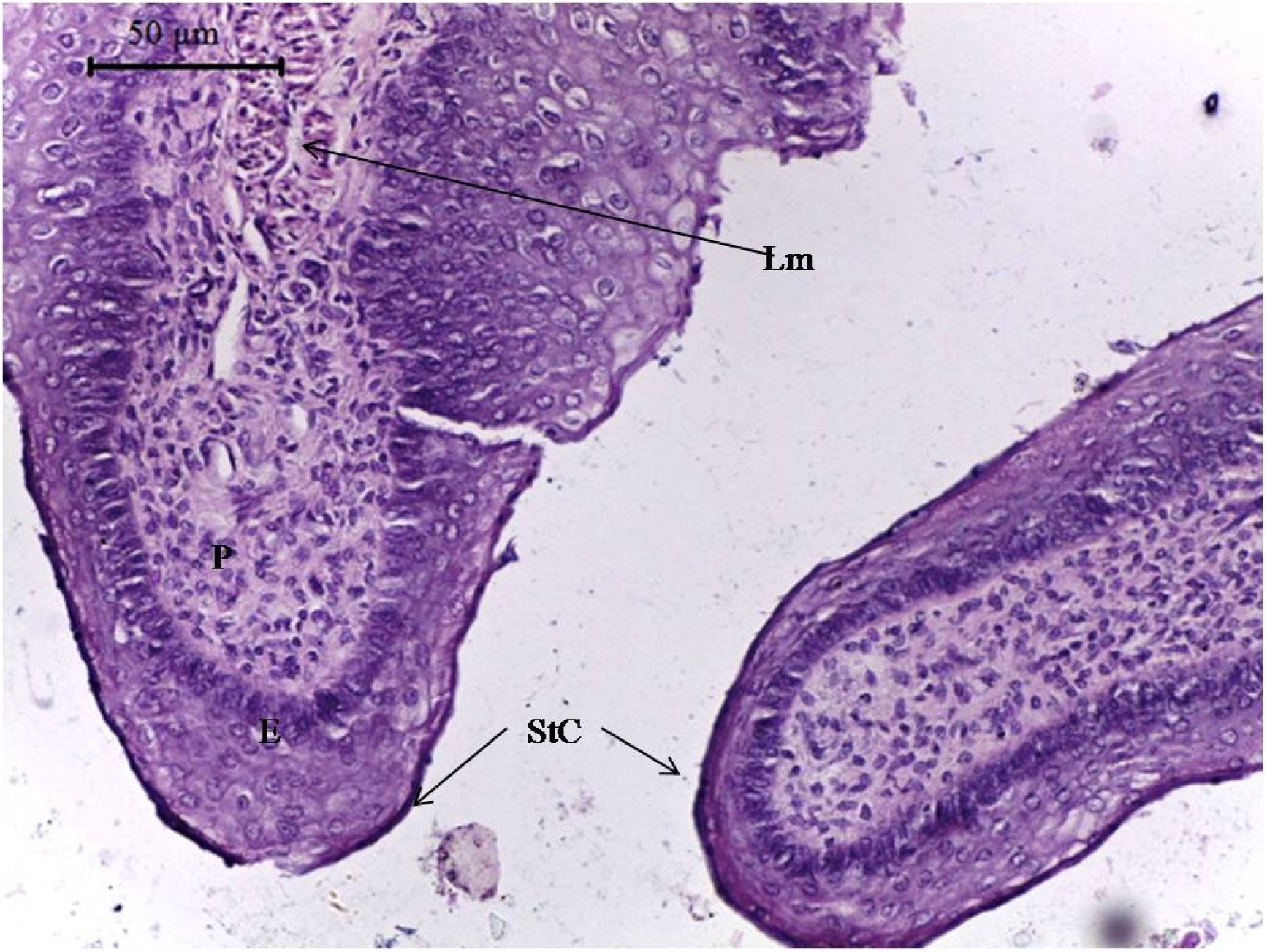
Photomicrograph of section of 145 day old goat foetal reticulum showing stratified squmous epithelium (E) of reticulum, stratum corneum (StC) and bundles of lamina muscularis (Lm) in the core of the primary reticular crest (P). H & E X 200

Wavy, pyramidal or dome shaped secondary reticular crests were noticed at 112 days of gestation. The tip of the secondary crest was rounded. The number of layers of superficial zone was more in secondary crest as compared to the primary reticular crest (Figure 6). At 121 days of gestation primary reticular crest showed lateral out pocketing, future corial papillae (Figure 6). Five to six cell layers of basal zone got congregated at certain level and formed wavy structure, corial papillae, which evaginated the superficial cell layers. These cells were columnar shaped with elongated nuclei and darkly stained eosinophilic cytoplasm. The number of corial papillae increased as age advanced. Corial papillae were also noticed in secondary reticular crest at about full term fetal reticulum (Figure 6). The primary reticular crest grew in two directions, longitudinal and transverse. Longitudinal growth as towards the epithelial surface followed by lateral growth to join with the adjacent rib resulted into formation of honey comb like structure.

Primary and secondary reticular crest and corial papillae contain all the contents of propria-submucosa. Vascularization increased with advancement of age. Few smooth muscle cells of tunica muscularis started migration towards the base of the primary reticular crest at 112 days of gestation. The propria submucosa contained sporadic differentiating smooth muscle cells along with above mentioned structures at 121 days of gestation. At 134 days of gestation migratory smooth muscle cells unite to form a layer below the basement membrane, fore runner of future lamina muscularis (Figure 7). Circularly arranged muscle bundles were recorded in proximal 1/3^rd^ of the primary reticular crest (Figure 9). Connective tissue cells and blood vessels were present in between the muscle bundles. The reticular fibers became coarser with advancement of age but were isolated type. At 121 days of gestation the reticular fibers joined with each other to form rete (Figure 5). In full term foetal reticulum bundles of short, thin to coarse loosely arranged reticular fibers were noticed in the lamina propria, whereas, in submucosa region these fibers were dense, intermingled with each other and arranged in regular bundles. The collagen fibers became mature from 121 days of gestation which were very thin and isolated type and were mostly noticed near or at the basement membrane (Figure 8). From 134 day onward few, isolated fine elastic fibers were observed in lamina propria and submucosa (Figure 10). Propria-submucosa was 53.31 ± 21.16 µm thick in this group.

**Figure 10.**
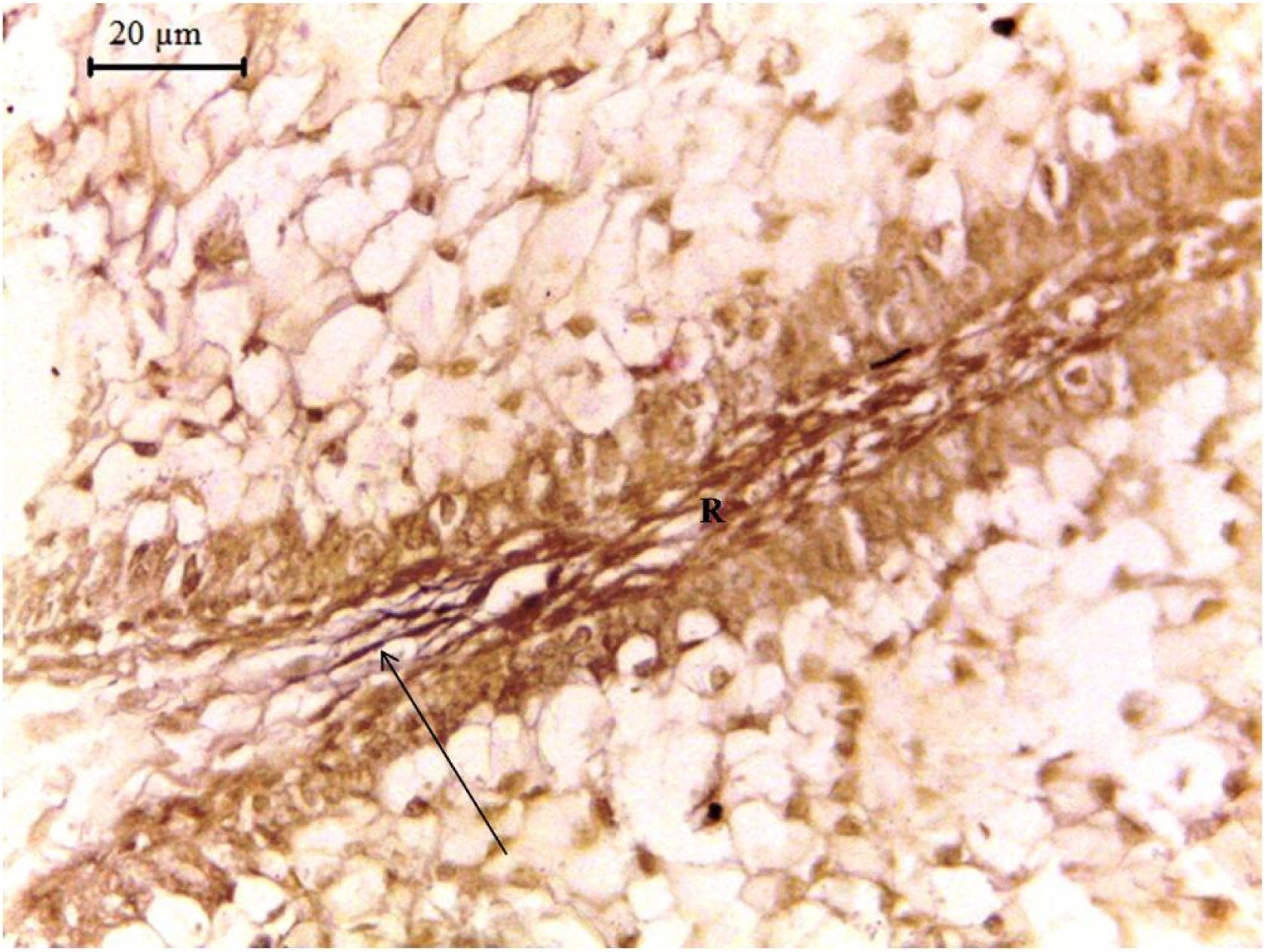
Photomicrograph of section of 134 day old goat foetal reticular wall showing elastic fibers in the core of reticular crest (R) (arrow). Verhoeff’s Stain X 400

At 112 days of gestation small clusters of smooth muscle cells were joined together and formed larger bundles. At 121 days of gestation circularly arranged bundles contained five to seven cells whereas, longitudinally directed bundles had three to four cells (Figure 6). Numerous coarse reticular fibers in rete form lied in between and within the smooth muscle bundles (Figure 5). Abundant collagen fibrils invaginating from submucosa to smooth muscle bundle were noticed at this stage. With the advancement of age both number and thickness of collagen fibers increased (Figure 8). Few isolated elastic fibers were also noticed surrounding the clusters of smooth muscle cells from 118 day onwards. Thickness of tunica muscularis and tunica serosa was 268.02 ± 131.69 and 15.54 ± 3.39 µm, respectively. From group II to group III thickness of epithelium, tunica muscularis and serosa increased while propria submucosa decreased.

## Discussion

At 32 days of gestation, the wall of the stomach consisted of three strata viz. epithelium, pleuripotent blastemic tissue and serosa. In contrary to this wall of the stomach differentiated into internal epithelium and external pluripotential blastemic tissue onlyin intrauterine life of sheepat 23-29 days [6], red deer 30 days [7] and goat 35 days [1]. Separate serosa layer was not mentioned in these animals. The present observations were in close agreement with the reports in sheep at 34 and 32 days of gestation, respectively [6, 8]; in goat at 38 and 28 days [1, 9] respectively; in cattle at 30 days [10] and in buffalo at 53 days of gestation [11]. At 32 days, the epithelium was undifferentiated stratified type, whereas, it was pseudostratified, non ciliated cylindrical stratified type in 23-29 day foetal sheep [6], 30 day foetal red deer [7] and stratified type of epithelium in goat at 35 day [1]. The pleuripotent blastemic tissue comprised of different shapes of mesenchymal cells with ground substance, blood vessels and immature red blood cells as reported earlier in sheep at 32 [6] and red deer at 30 [7] days of gestation. The wall of the stomach was surrounded by a single layer of squmous cells, the mesothelium.

Microscopically the reticulum became independent compartment at 38 days of gestation. At this stage it was in the form of compact cell mass with canalization at the center. In goat differentiation of reticulum was reported at 35 days [12, 13], while in sheep and red deer 33 and 60 days, respectively [14, 15]. The reticulum gets differentiated in buffalo foeti at 3.5 cm Crown Rump Length (CRL) [16].

The epithelium was undifferentiated stratified type. From 38 days onwards the epithelium showed 2 distinct zones viz. darkly stained basal and lightly stained superficial zone. An increase in the number of layers of basal and superficial zone was noticed upto 59 and 100 days of gestation, respectively and thereafter decreasing trend was observed. With the advancement of age cytological characters of cells of deepest layer remained constant however, their cytoplasm became more eosinophilic and foamy near the term. The cytological characters of basal zone were in close proximity with the earlier reports in buffalo [16, 17]. At 87 days few of the cells of middle layer showed characters of future stratum spinosum. At term few cells of topmost layer had lost their nuclei. The cytoplasm of those cells was highly eosinophilic and referred as stratum corneum. In addition to stratum spinosum; stratum granulosum and stratum lucidum were also recorded in sheep foetal reticulum during 81-112 days of gestation [14]. However, these authors mentioned that the cells of stratum lucidum/ spinosum were large in size but poorly defined while the cells of stratum corneum were elongated anuclear cells arranged parallel to the surface. They further reported that the epithelium remained non-keratinized till birth. The beginning of formation of stratum corneum in the present study was in proximity of earlier reports in sheep at 53 and at birth, respectively [8, 18]. Stratum corneum was noticed at 104 and 142 days of gestation in goat and red deer [9, 11]. Keratinization of reticular epithelium was reported in red deer foeti at 205 days of gestation and buffalo at 74 cm CRL [15, 16].

The first sign of reticular crest formation was noticed at 60 day of gestation with evagination of basal zone epithelium and simultaneous invagination of content of only lamina propria as reported earlier in buffalo at 10.2 cm CRL [16] and goat at 59 days of gestation [12, 13]. The reticular crest was observed as evagination of basal zone of epithelium into superficial zone at 16 cm CRL Egyptian water buffalo [19]. These crests were also referred as reticular cellulae and were reported at 64 days of gestation in sheep foeti as a simple evagination of stratum germinativum. These cellulae became more prominent at 69 days of gestation and it involved only lamina propria without affecting submucosa [14]. In bovine foetal reticulum primary reticular crest formed between 50-60 days [20] and 3^rd^ month of gestation [21], whereas, secondary crests were observedat 95 days [19]. The process of reticular crest formation was further progressed with the advancement of age. The shape of the crest varied from crecenteric to low pyramidal or pyramidal from 70-87 days of gestation. Pyramid shaped reticular crests were also noticed in buffalo foetal reticulum [16]. The height of the primary reticular crest increased with the advancement of age. Inter reticular crest distance of primary reticular crest increased as age advanced due to formation of secondary reticular crest between them. Dome shaped secondary reticular crest were first observed at 112 days of gestation. Height of this crest was always lower than the primary reticular crest. Appearance of secondary reticular crest was also observed in buffalo at 25.5 cm CRL [16] and in sheep at 30cm and 26 cm CRL [8, 22]. The core of the primary reticular crest in its proximal 1/3^rd^ part contained circularly oriented smooth muscle bundles, muscularis mucosae near term. Muscularis mucosae when contract reduces the height of the crest [23]. It was suggested that contraction of these smooth muscle reduced the epithelial height, facilitated the movement of blood away from primary fold after its vessels had became filled during digestion, absorption of Volatile fatty acid, ammonia, sodium, potassium etc. The connective tissue elements were more at the base and towards the tip. These observations were in accordance with the 50 cm CRL Egyptian water buffalo foetus [19] at 113 days of gestation in sheep [14], in 50.5 and 60.1 CRL buffalo foetus [16, 17] and 101 days in gestation in goat [13]. At 121 days of gestation, corial papillae were observed as lateral out pocketing of primary reticular crest and their number increased with the advancement of age. These corial papillae were also noticed in the secondary reticular crest near the term. The core of the corial papillae contained lamina propria only. The present study was in partial agreement withgoat foetal reticulum corial papillae located on the lateral surface of primary reticular crest at 101 days referred as corneum papillae [13] and corial papillae between 81-112 days of gestation in sheep [14]. In buffalo foeti corial papillae were recorded at 20 cm CRL [17]. However, corial papillae could not detect till birth in sheep and goat foetal reticulum [8, 9].

Lamina propria, submucosa and tunica muscularis were represented by pleuripotent blastemic tissue up-to 49 days of gestation. Blastemic tissue was composed of differentiating mesenchymal cells, differentiating fibroblasts, differentiating myocytes, capillaries, few neuronal elements and ground substance. Fine reticular fibrils were present in the blastemic tissue around differentiating myocytes and blood vessels. The reticular fibers became coarser at 100 days of gestation and these fibers formed network at 121 days of gestation. Reticular fibers were more in submucosa region and interwoven with adjacent fibers as reported in 14.7-19.6 cm CRL buffalo foeti [17]. Immature, isolated collagen fibers observed in propria submucosa at 100 days which became coarser at 121 days of gestation. The elastic fibers were observed in propria submucosa and large blood vessels at 134 days of gestation. The appearance of elastic fibers in the propria submucosa was in harmony with the findings in sheep (120 days of gestation) and full term buffalo foetus [14, 17].

At 51 day of gestation there was no clear separation between propria and submucosa as reported by in sheep [14, 18] and buffalo [16, 17]. Lamina propria was densely populated with mesenchymal cells and darkly stained while submucosa was less cellular with profound ground substance and lightly stained. Few myocytes were arranged in clusters below the propria submucosa forming a distinct tunica muscularis similar to the observation in sheep at 52 days of gestation in [22], at 67 days of intra uterine life in red deer [7] and from 3.2 cm and 5.5 cm CRL in buffalo foeti [17, 24]. On the contrary early appearance of tunica muscularis was also noticed at 33 and 34 days of gestation in sheep [14, 25]. At 112 days of gestation few smooth muscle cells started migration towards the base of reticular crest i.e. future lamina muscularis. At 134 days of gestation distinct layer of lamina muscularis mucosae was evident. Similar findings had been reported in primary reticular rib at 113 days of gestation in sheep [14].

In the present study orientation of smooth muscle layers was not constant. At locations tunica muscularis was comprised of three layers viz. circular, longitudinal and few obliquely directed smooth muscle fibers. The reason attributed to the reverse orientation of muscle fibers in the wall of reticulum was possibly due to spiral and oblique arrangement of muscle fibers. Similar observations were recorded in goat [26] and buffalo foeti [17]. At term the muscle fibers were arranged longitudinally towards the propria submucosa and circularlytowards the serosa. A progressive increase in thickness of tunica muscularis with increase of foetal age was recorded as described previously in buffalo [17]. There was about three times increase in thickness from group II and III. Neuronal elements were noticed in between the clusters of myocytes.

At 38 days of gestation developing reticular wall was lined by a single layer of differentiating mesothelial **c**ell. From 51 days onwards this layer was well supported by loose vascular connective tissue and had profuse amount of ground substance as reported in bovine at 53 days of gestation [20]. Insheep and buffalo foetal reticulum mesothelial lining was well supported by loose connective tissue and as age advanced it was having well developed connective tissue elements, large blood vessels and neuronal elements [14, 17]. Thickness of tunica serosa increased with advancement of age. In contrary to this decreasing trend was noticed in sheep [14].

## Materials and Methods

The present study was conducted on the developing reticulum collected from 36 healthy and normal embryos/ foeti of either sex of non-descript goat (*Capra hircus*) of Mathura region of India. An approval was obtained from animal ethic committee of DUVASU, Mathura (U.P.)India prior to the commencement of the study.

### Collection of material

The embryos/ foeti were ranged from 32 days to near full term (145 days). The age of embryos/foeti was ascertained by using formula for goat foetus, W^1/3^ = 0.096 (t-30), where W = body weight of foetus in gram and t = age of foetus in days [3]. Embryos/foeti were assigned into three group viz. group I (0-50 days of gestation), group II (51-100 days of gestation) and group III (101-150 days of gestation). The abdominal cavity was opened and developing reticulum was harvested. Small pieces of tissues were cut in group II and III while in group I whole of the stomach was collected. The tissues were fixed in 10 per cent neutral buffered formalin and were processed by routine paraffin embedding technique.

### Staining techniques

6 μm thick sections were taken and stained with hematoxylin and eosin for histo architecture, Wilder’s reticulin stain for reticular fibres [4], Verhoeff’s stain for elastic fibres [4] and Mallory’s triple stain [4] for collagen fibers. Stained slides were observed under light microscope. Micrometric observations were done on hematoxyline eosin stained sections by Leica DM750 computerized image.

### Statistical analysis

The data generated by the micrometrical observations were subjected tostatistical analysis [5] with the help of SPSS 20.0 software.

